# Evolutionary trajectories of resistant mutants during sub-MIC antibiotic exposure

**DOI:** 10.1101/2024.05.25.595866

**Authors:** Omar M. Warsi

**Author notes:** ***Conflict of Interest:*** There are no competing interests to be declared.

## Abstract

The emergence of antibiotic resistance is one of the most important examples of contemporary evolution. Selection for resistance can occur over a wide concentration range, both above and below the minimum inhibitory concentration (MIC) of the antibiotic. In a majority of cases, resistance mutations confer fitness costs and several studies have shown the importance of these costs for the emergence, ascendance and maintenance of resistance in a population. Importantly, these costs can often be ameliorated by compensatory mutations and rate and efficiency of compensation is a key parameter in determining the evolutionary success of a costly resistance mutation. Despite this knowledge, we still have a limited understanding of how resistance evolution (to increase resistance) and compensatory evolution (to reduce fitness costs) interact during growth in presence of low, sub-MIC, antibiotic concentrations. To examine the impact of these two processes, we carried out evolution experiments at sub-MIC levels of streptomycin using two *E. coli* mutants (with loss of function mutations in the *selB* and *ubiH* genes, respectively) that show low-level streptomycin resistance, and have different fitness costs. For both mutants, evolution at sub-MIC levels enriched for mutations that increased resistance, but selection for compensatory mutations was also common over the course of the experiment. Our study highlights that costly low-level resistant mutants adapt to sub-MIC antibiotic exposure by either increasing resistance, reducing cost or both and that this evolution can result in the maintenance of these mutants in the population.

## INTRODUCTION

The use of antibiotics during the last 80 years has selected for resistant bacteria that are more difficult to treat, resulting in increased morbidity and mortality (Levy and Marshall 2004; Ferri et al. 2017; Naylor et al. 2019). To reduce and predict the emergence of resistant bacteria, there is a need to understand the mechanistic basis and dynamics of selection of antibiotic resistance. Antibiotic resistance evolution in a given environment is a function of the concentration of the antibiotic present in the environment (Dahlberg and Chao 2003; Bergman et al. 2004; Olofsson et al. 2007; Bergman et al. 2009). At high-antibiotic concentrations that are above the minimum inhibitory concentration (MIC) of the antibiotic, high- and low-cost resistance mutations can get selected and be maintained in the population since the susceptible bacteria would not survive at these concentrations (Fig.1A). On the other hand, evolution of resistance at sub-MIC of antibiotics is expected to result in selection of only low-cost resistance mutations, since selection at sub-MIC is weak and any resistance mutations that have a cost higher than the selective effect of the antibiotic could therefore not rise in frequency in the population (Fig.1B) (Gullberg et al. 2011; Wistrand-Yuen et al. 2018). This difference in resistance mutation spectra based on the concentration of the antibiotic in the environment has been corroborated by several studies (Westhoff et al. 2017; Wistrand-Yuen et al. 2018; Sanz-García et al. 2020), highlighting the need to understand how and why the mechanisms of antibiotic resistance evolution at antibiotic concentrations above and below the MIC differ.

**Fig. 1.**
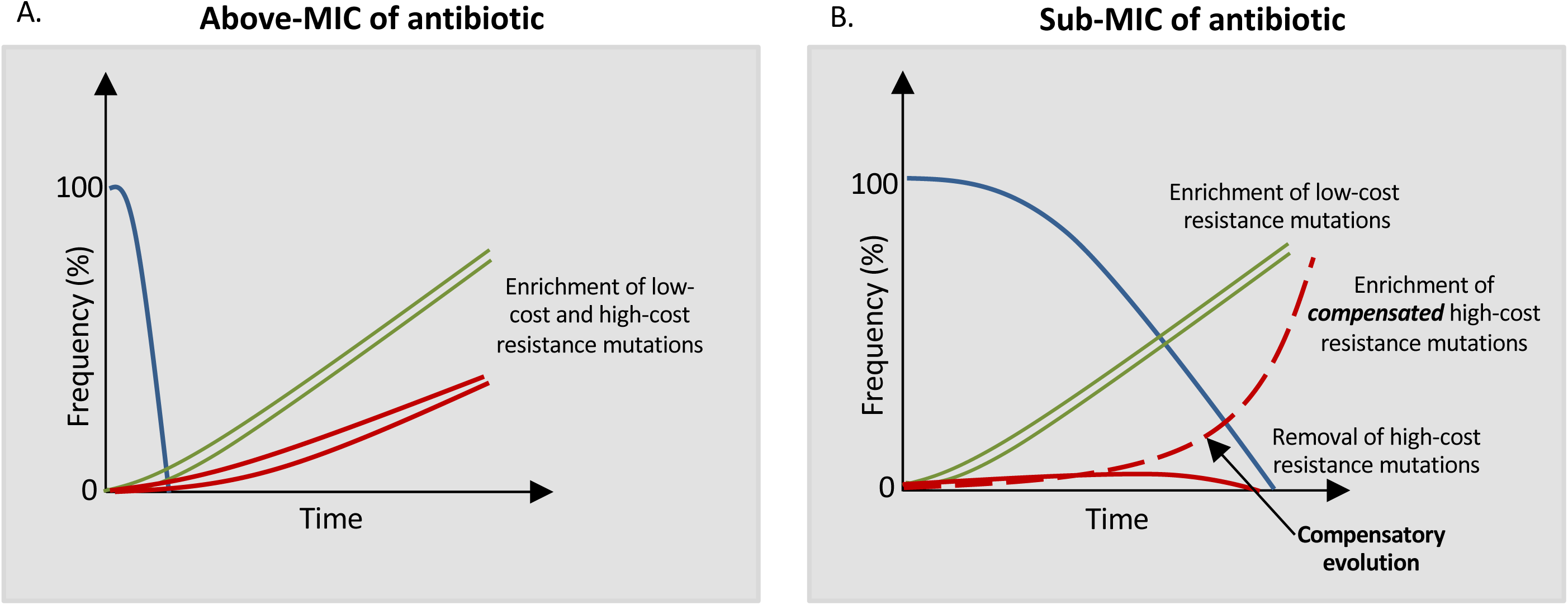
Characteristics of antibiotic resistance evolution at above- and sub-MIC of antibiotic. **A)** Trajectories of high-cost (red) and low-cost (green) resistant mutants depicting the enrichment of these resistant mutants at above-MIC of antibiotic. At these concentrations, the susceptible ancestral bacteria (blue) are quickly outcompeted by both types of resistant mutants. **B)** Susceptible ancestral bacteria (blue) are outcompeted by the low-cost (green) resistant mutants at sub-MIC of an antibiotic. High-cost resistant mutants (red), in most cases, will get outcompeted by both the wild-type and the low-cost resistant mutants. However, some high-cost resistant mutants can also get enriched at sub-MIC antibiotic levels if compensatory evolution is sufficiently fast and efficient (dotted red-line).

Understanding the evolutionary processes that result in enrichment of resistant mutants at sub-MIC of antibiotics is important since low concentrations of antibiotics are found in numerous different environments, including soil, surface water, wastewater treatment plants and in antibiotic-treated humans and animals. In these environments, low-level resistance mutations, given their high frequency, would generally be the first step for evolution of high-level resistance (Wistrand-Yuen et al. 2018). However, the evolutionary trajectories that these low-level resistant mutants would undertake at continued sub-MIC antibiotic exposure with regard to resistance level and compensatory evolution is unclear, and is expected to depend on both the fitness cost and the rate and efficiency of subsequent compensatory evolution (Fig.1B).

Most studies of compensatory evolution of resistant mutants with reduced fitness has been done in absence of antibiotics and demonstrated that compensatory mutations generally increase fitness of the resistant mutant by either directly or indirectly correcting the physiological defect caused by the resistance mutation (Bjorkman et al. 2000; Andersson and Hughes 2010; Qi et al. 2016; Brandis and Hughes 2018). However, at sub-MIC of an antibiotic continuous selection for antibiotic resistance might result in some of these compensatory mutations being inaccessible since reversion of the resistant phenotype to susceptibility will also result in decrease in fitness. Alternatively, continuous selection for increasing antibiotic resistance might result in new routes to increase fitness, without necessarily involving a correction of the physiological defect caused by the initial resistance mutation. Studies focused on selecting resistance mutations at sub-MIC of antibiotics show that there is a continuous selection for increasing resistance in these populations (Gullberg et al. 2011; Gullberg et al. 2014; Westhoff et al. 2017; Wistrand-Yuen et al. 2018); however in only one study the role of compensatory mutations has been inferred (Westhoff et al. 2017).

To examine how antibiotic resistance evolution and compensatory evolution interacts under weak selective pressures of sub-MIC of antibiotics, we first isolated low-level streptomycin resistant mutants of *E. coli* and then evolved them further at sub-MIC of streptomycin to analyze their evolutionary trajectories. Our experimental design consisted of growing two low-level resistant mutants with different fitness costs at sub-MIC concentrations where the selection for resistance mutations would be strong (close to the MIC of the ancestral susceptible strain) and where the potential for compensatory evolution would vary because the mutants had different fitness costs. This allowed us to determine if, apart from selection for increased resistance, compensatory evolution occurred at sub-MIC antibiotic levels, and how the fitness cost of the resistance mutation would affect the process. Our findings show that selection for increased resistance was the dominant outcome, but for both mutants compensatory evolution also occurred, suggesting that there is concomitant selection for both outcomes during sub-MIC exposure.

## MATERIALS and METHODS

### Bacterial strains, growth media and MIC determination

The bacterial strain *Escherichia coli* K-12 MG1655 was used in all experiments. The streptomycin resistant mutants derived from this ancestral strain are listed in Table 1. The nutrient rich media used were Lysogeny Broth (LB) (liquid media) and LB Agar (solid media, Sigma-Aldrich). All experiments were performed at 37°C. Minimum inhibitory concentrations (MIC) on solid media were determined by plating 10^-7^ dilution of an overnight grown culture on different concentrations of streptomycin (2-fold dilution steps). Using this method, the MIC of streptomycin for the wild-type strain on solid media was determined to be 4 mg/L, and this is designated as MIC_wt_solid_ throughout the manuscript. MIC determination was also performed for this strain in liquid media, and was determined to be 10 mg/L (Supplementary figure 1), and is designated as the MIC_wt_liquid_ throughout the manuscript.

**Table 1:**
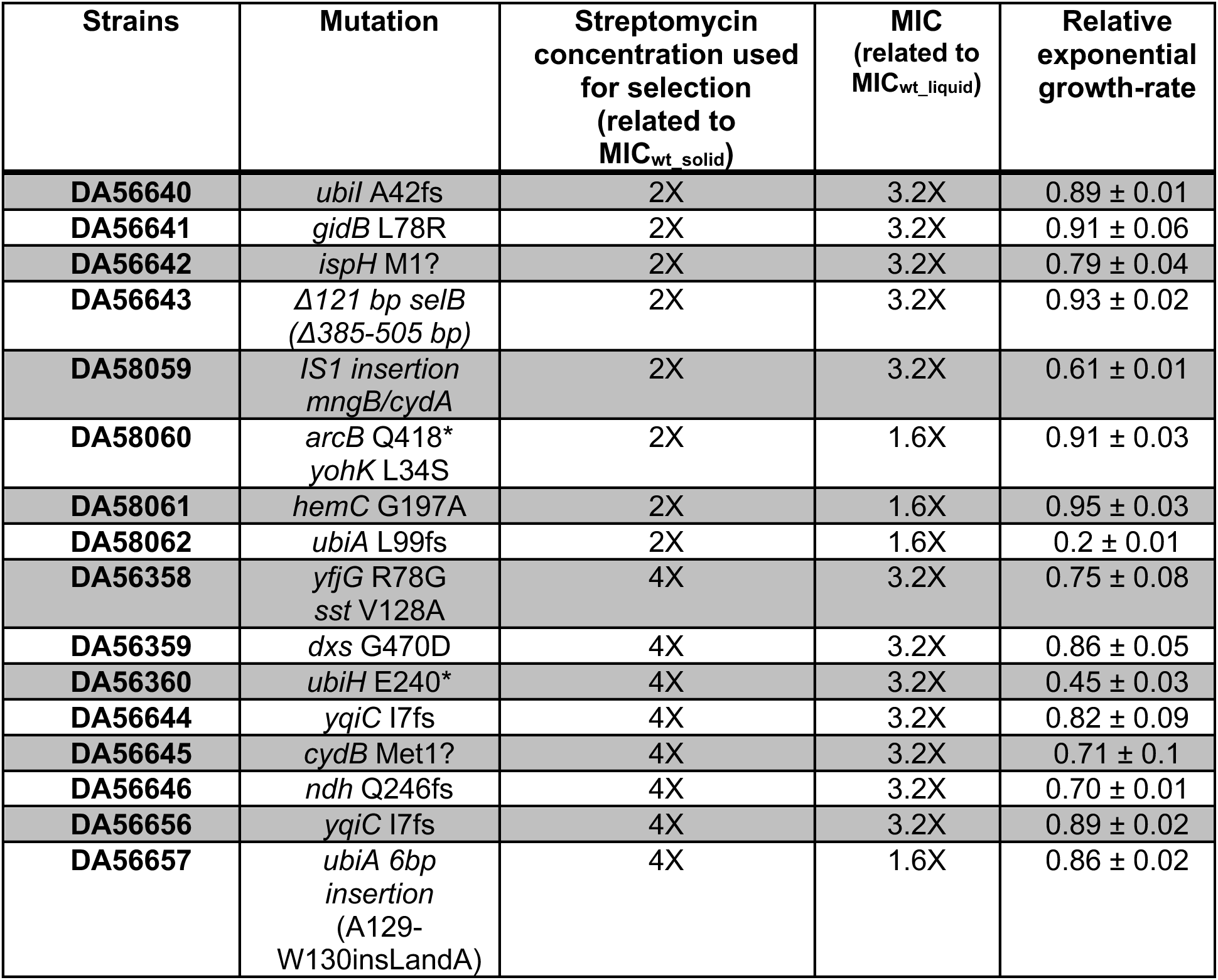
Low-level streptomycin resistant *E. coli* mutants selected at 2X MIC_wt_solid_ and 4X MIC_wt_solid_ streptomycin concentrations (MIC_wt_solid_ = 4mg/L). Growth rates are shown relative to growth rate of susceptible MG1655 strain measured in the absence of the antibiotic. All amino acids are denoted as single letter codes. A * denotes a stop-codon, adenotes a mutation in start-codon, and a fs denotes a frame-shift mutation.

| Strains | Mutation | Streptomycin concentration used for selection (related to MIC<sub>wt_solid</sub>) | MIC (related to MIC<sub>wt_liquid</sub>) | Relative exponential growth-rate |
| --- | --- | --- | --- | --- |
| DA56640 | *ubiI* A42fs | 2X | 3.2X | 0.89 ± 0.01 |
| DA56641 | *gidB* L78R | 2X | 3.2X | 0.91 ± 0.06 |
| DA56642 | *ispH* M1? | 2X | 3.2X | 0.79 ± 0.04 |
| DA56643 | $\Delta 121$ bp *selB* ($\Delta 385$-505 bp) | 2X | 3.2X | 0.93 ± 0.02 |
| DA58059 | *IS1* insertion *mngB/cydA* | 2X | 3.2X | 0.61 ± 0.01 |
| DA58060 | *arcB* Q418*<br>*yohK* L34S | 2X | 1.6X | 0.91 ± 0.03 |
| DA58061 | *hemC* G197A | 2X | 1.6X | 0.95 ± 0.03 |
| DA58062 | *ubiA* L99fs | 2X | 1.6X | 0.2 ± 0.01 |
| DA56358 | *yfjG* R78G<br>*sst* V128A | 4X | 3.2X | 0.75 ± 0.08 |
| DA56359 | *dxs* G470D | 4X | 3.2X | 0.86 ± 0.05 |
| DA56360 | *ubiH* E240* | 4X | 3.2X | 0.45 ± 0.03 |
| DA56644 | *yqiC* I7fs | 4X | 3.2X | 0.82 ± 0.09 |
| DA56645 | *cydB* Met1? | 4X | 3.2X | 0.71 ± 0.1 |
| DA56646 | *ndh* Q246fs | 4X | 3.2X | 0.70 ± 0.01 |
| DA56656 | *yqiC* I7fs | 4X | 3.2X | 0.89 ± 0.02 |
| DA56657 | *ubiA* 6bp insertion (A129-W130insLandA) | 4X | 1.6X | 0.86 ± 0.02 |

### Isolation of low-level resistant mutants and calculation of mutation rate

The mutation rate of resistance at different streptomycin concentrations was calculated by a fluctuation test. Briefly, ∼10^3^ cells from an overnight culture were used to start 40 independent cultures. From these overnight grown cultures, ∼10^6^ cells were plated on plates containing streptomycin concentrations of either 2X MIC_wt_solid_ or 4X MIC_wt_solid_. The mutation rate was determined using the frequency of resistant mutants and the bz-mutation rate calculator (Gillet-Markowska et al. 2015).

### Whole genome sequencing and sequence analysis

To identify resistance mutations and other adaptive mutations in the evolving populations, single clones were isolated and whole genome sequenced. DNA extraction was done with 1 ml of overnight grown cultures using the Epicentre DNA extraction kit following the manufacturer’s protocol. Illumina’s Nextera XT kit was used to make whole-genome DNA libraries (2 x 300) that were then sequenced on Illumina’s Miseq platform. Samples were dual-indexed and pooled together. Average whole genome coverage per sample was 30X. Analysis of the fastq files obtained from Miseq sequencing was performed using CLC genomics Workbench version 8. This tool was used to identify point mutations and small indels. To identify large deletions or transposon movements, the fastq files were also analyzed using Breseq (version 0.27.1a) (Deatherage and Barrick 2014).

### Growth rate measurements and calculation of fitness costs

To measure the fitness of evolved populations and clones, exponential growth rate measurements were performed using a BioscreenC analyzer at OD_600_, with measurements taken every 4 minutes. KaleidaGraph software was used to calculate the maximum exponential growth rate from the OD_600_ data, using the optical density (OD) values between 0.02 to 0.09. Fitness costs were calculated as the ratio of exponential growth rate for the resistant mutant to that of the wild-type strain, measured in the absence of the antibiotic. Similarly, growth-rates of different mutants at different streptomycin concentrations were plotted as a ratio between exponential growth rate of the mutant at a given streptomycin concentration to that of the wild-type strain in absence of any antibiotic. Growth-rates of evolved clones of mutants *ubiH* and *selB* were represented as the ratio between exponential growth rate of the evolved clone at a given streptomycin concentration to that of the ancestral mutant strain in absence of any antibiotic. As before, for each strain three biological replicates were used, and the values in the figures represent the average of the biological replicates while the error bar represents the standard deviation. For calculation of population exponential growth rates, six replicates were used in each case.

### Evolution experiments and screening for high-level resistant mutants

Evolution experiments were carried out in batch cultures, where ten independent populations for each mutant were grown overnight in 1 ml LB medium containing 8mg/L of streptomycin (0.8X MIC_wt_liquid_). The next day, 1μl of the culture was transferred into 1 ml fresh LB medium and allowed to grow again for 24 hrs, meaning that each population grew for ∼10 generations every day. This was done for a period of 25 days, approximating ∼250 generations. For each population, populations were saved at −80°C in 10% DMSO after every 50 generations. To determine the frequency of high-level resistant mutants in each of the evolving populations, 1ul (∼10^6^) cells were plated on LA plates containing either 64 mg/L (16X MIC_wt_solid_) or 100 mg/L (25X MIC_wt_solid_) streptomycin. To determine total population density that was being plated, dilutions of each population were also plated on plates without antibiotic. Fixation of high-level resistant mutants was assumed when a bacterial lawn was observed on plates containing either 64 mg/L (16X MIC_wt_solid_) or 100 mg/L (25X MIC_wt_solid_) streptomycin.

### Construction of different rpsL variants

The different *rpsL* mutants were constructed using the λ-red recombineering technique (Yu et al. 2000). This involved using 500 ul of an overnight grown culture of *E. coli* K-12 MG1655 strain containing the pSIM5-Tet plasmid (Koskiniemi et al. 2011) (grown at 30°C) and inoculating 50 ml of LB broth containing 10 mg/L of tetracycline. These cells were allowed to grow at 30°C until the optical density (OD 600) reached 0.2, after which they were transferred to a 42°C shaking water bath for 20 mins. At the end of 20 mins, the cells were placed on ice for 5 minutes and were then washed thrice with 10% glycerol. These cells were then transformed by electroporation using single-stranded oligonucleotides (100 nucleotides in length) that contained the desired *rpsL* mutation. After transformation, the cells were recovered overnight, and were plated on streptomycin-containing plates (25 mg/L or 100 mg/L). Transformants were re-streaked and the *rpsL* gene was PCR-amplified and Sanger sequenced to confirm that the desired mutation was inserted.

### Competition experiments between the ubiH mutant and the susceptible ancestral E. coli strain

Competition experiments were performed by genetically tagging the different strains with genes encoding for fluorescent markers (yellow fluorescent protein (YFP) and blue fluorescent protein (BFP)). Fluorescently labeled strains were grown over-night in the absence of streptomycin, and then mixed in a 1:1 ratio to start the competition experiments. All competitions were performed in 96-well plates, where 2 μl of an overnight-grown culture was transferred to 200 μl of media each day, resulting in ∼6.6 generations per day. The frequencies of each strain were determined at five time-points (0, ∼13, ∼30, ∼50 and ∼100 generations) by making 100-fold dilutions of cultures from each well in phosphate buffered saline for flow cytometry. 100,000 cells were counted and the fraction of BFP and YFP positive cells was determined by flow cytometry (MACSQuant VYB, Miltenyi Biotec). The rate of change of the different strains is plotted as the natural log of the ratio of different strains at different time-points.

## RESULTS

### Distribution of fitness effects of low-level streptomycin resistant mutants at sub-MIC of streptomycin

We selected streptomycin resistant mutants of the wild-type *E. coli* K-12 strain MG1655 (MIC of wild-type *E. coli* K-12 on solid agar media, henceforth designated as MIC_wt_solid_ = 4 mg/L) at concentrations two times (2X MIC_wt_solid_ = 8 mg/L) and four times (4X MIC_wt_solid_ = 16 mg/L) the MIC of streptomycin by growing 40 independent over-night cultures of the susceptible bacteria and spreading them on independent LB agar plates containing the two different concentrations of streptomycin. The mutation rate for streptomycin resistance was calculated to 3.3 x 10^-7^ and 4.2 x 10^-7^ at 2X and 4X MIC_wt_solid_, respectively. We isolated 16 independent mutants by randomly picking one colony from the selection plates, and then whole-genome sequenced them to identify the mutations conferring the resistance phenotype (Table 1). We then determined the fitness effects of these resistance mutations both in the absence of the antibiotic, as well as at sub-MIC of the antibiotic in liquid (Fig. 2). The MIC of the wild-type susceptible *E. coli* strain in liquid media (MIC_wt_liquid_) was 10 mg/L (Supplementary figure 1), and all the growth measurements in liquid culture were based on this value. The general trend observed from these measurements was that all mutants had fitness costs in the absence of the antibiotic and also displayed a reduction of growth rate as the concentration of the antibiotic was increased (Fig. 2).

**Fig. 2.**
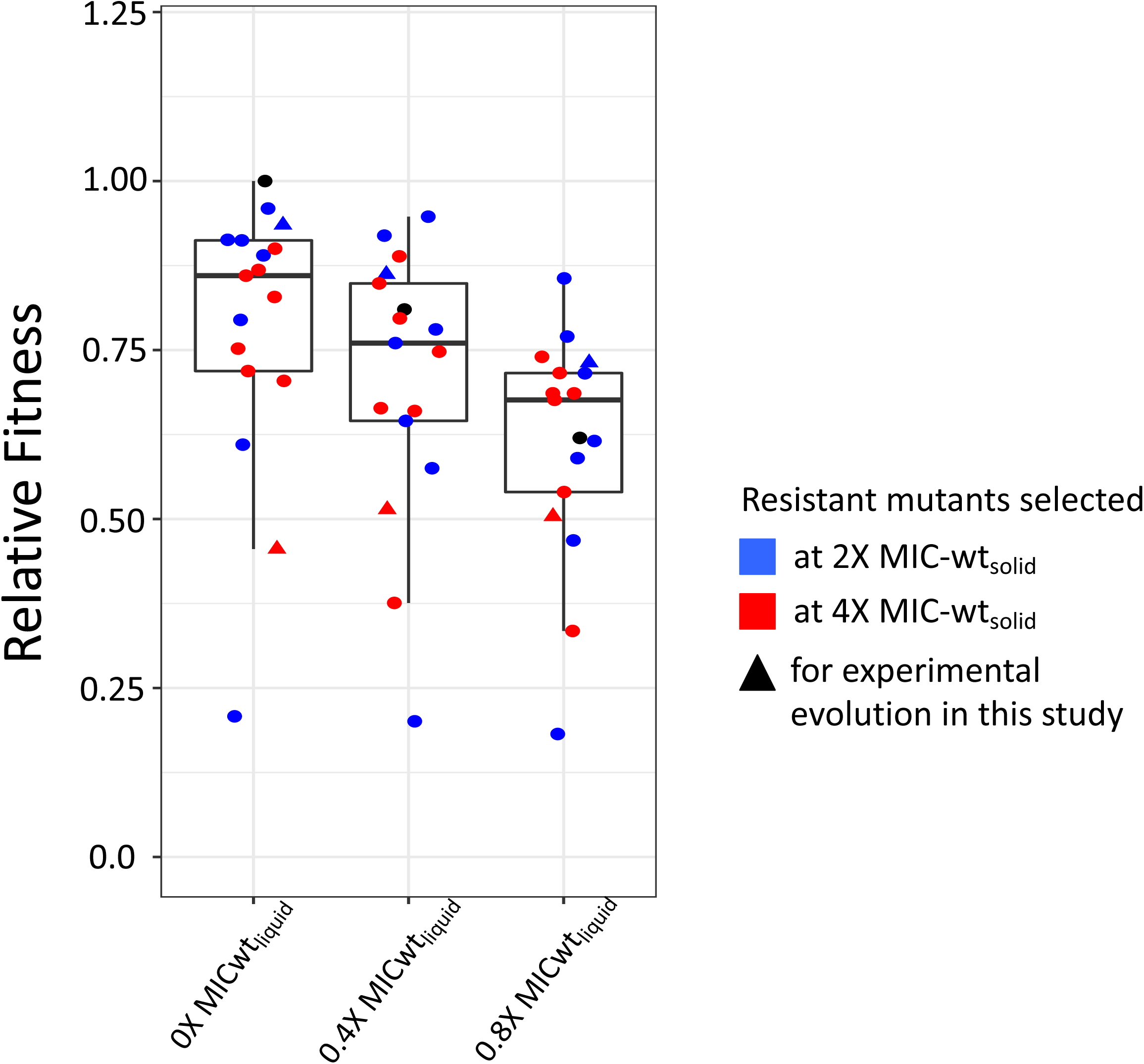
Distribution of fitness effects of streptomycin resistant *E. coli* mutants selected at streptomycin concentrations of 2X MIC_wt_solid_ and 4X MIC_wt_solid_. Relative growth rates of streptomycin resistant mutants selected at streptomycin concentration of 2X MIC_wt_solid_ (blue circles) and 4X MIC_wt_solid_ (red circles) were measured at different concentrations of streptomycin (no antibiotic, 0.4X MIC_wt_liquid_, and 0.8X MIC_wt_liquid_). Growth rates are shown relative to growth rate of susceptible MG1655 strain measured in the absence of the antibiotic. Three biological replicates were used in each case. Error bars show standard deviations. Growth of the susceptible ancestral MG1655 strain at each concentration is represented as black circles, while growth rates of mutants selected for evolution experiments are shown as triangles.

We chose two mutants for the analysis of compensatory evolution at sub-MIC of streptomycin. These mutants had mutations in the gene *selB* (Δ 121bp) and *ubiH* (E240*) (premature stop codon at position 240), respectively, and identical MICs of 3.2X MIC_wt_liquid._ The *selB* gene codes for a selenocystiene tRNA-specific translation elongation factor, while *ubiH* codes for an enzyme involved in ubiquinone biosynthesis. These mutants were chosen because they had different fitness costs in the absence of the antibiotic, and displayed different trends of changes in growth-rate at increasing streptomycin concentrations (Fig. 3). The *selB* mutant had a fitness cost of ∼7% in the absence of the antibiotic, and at increasing streptomycin concentrations the growth-rate was further reduced (Fig. 3). The *ubiH* mutant had a fitness cost of ∼50% in the absence of the antibiotic but increasing streptomycin concentrations had no observable effect on the growth rate (Fig. 3).

**Fig. 3.**
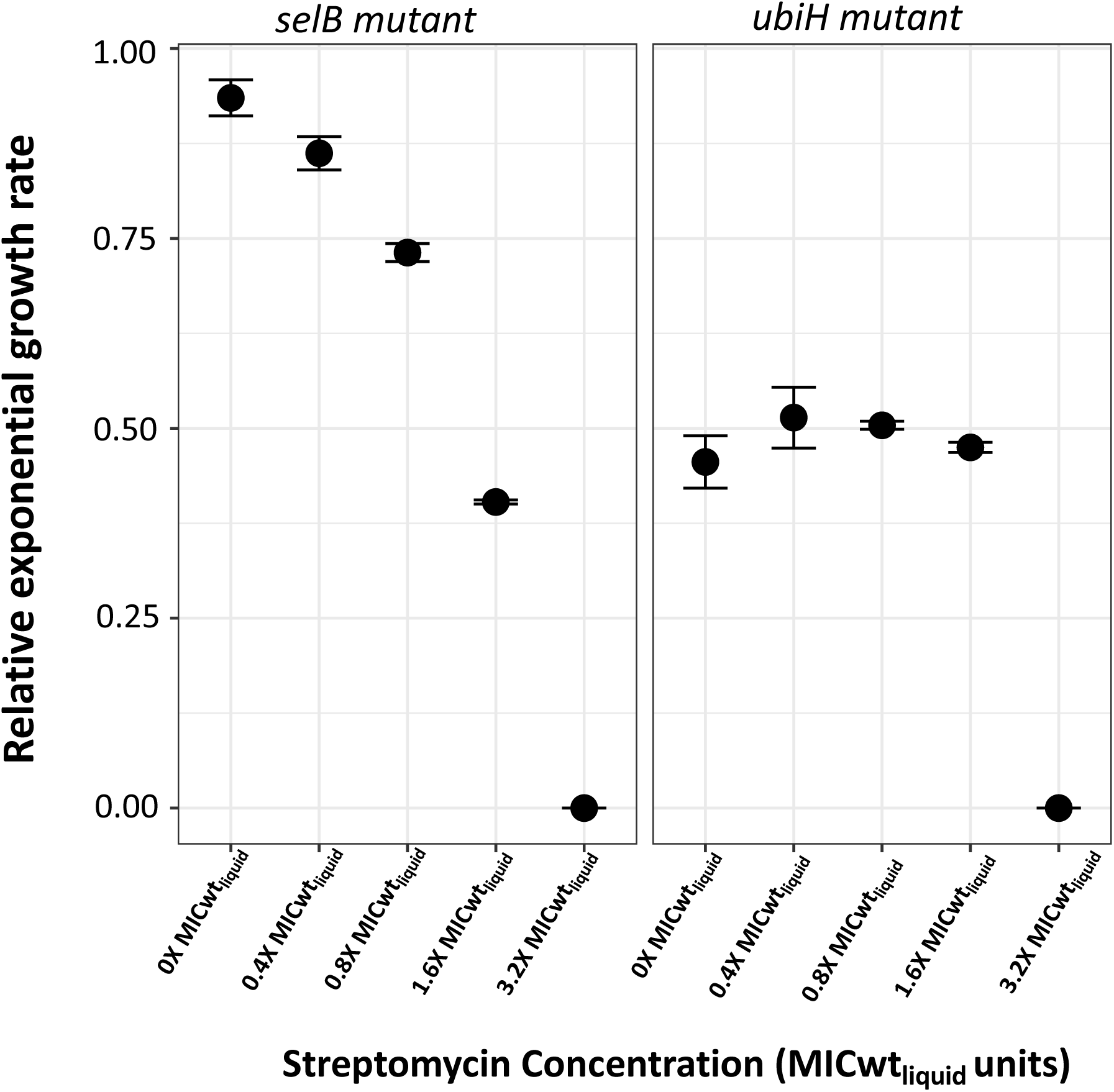
Effect of different concentrations of streptomycin on growth rates of the *E. coli selB* and *ubiH* mutants. Relative growth rates of *selB* and *ubiH* mutants at different streptomycin concentrations (no antibiotic, 0.4X MIC_wt_liquid_, 0.8X MIC_wt_liquid_, 1.6X MIC_wt_liquid_, and 3.2X MIC_wt_liquid_) was measured. Growth rates are shown relative to growth rate of susceptible MG1655 strain measured in the absence of the antibiotic. Three biological replicates were used in each case. Error bars show standard deviations.

### Evolution of *ubiH* and *selB* mutants at sub-MIC levels of streptomycin

Given that the MIC of streptomycin for the susceptible ancestral *E. coli* strain in liquid media was 10 mg/L, the compensatory evolution experiments were carried out at sub-MIC levels just below 10 mg/L. Thus, we serially passaged ten independent populations of each mutant (*ubiH* and *selB)* at 8 mg/L (i.e. 0.8X MIC_wt_liquid_). The MIC of streptomycin for these mutants was 32 mg/L or 3.2X MIC_wt_liquid_ at the start of the experiment. After 250 generations of growth, relative growth rates of each of the ten populations were measured with and without the antibiotic (Fig.4). The former was done to determine the increase in fitness of the populations at sub-MIC of streptomycin, while the latter was done to determine the level of compensatory evolution (or media adaptation) that might have taken place in these populations during the course of the experiment. All twenty populations (ten for each mutant) showed an increase in fitness at a streptomycin concentration of 8 mg/L (0.8X MIC_wt_liquid_), while 12 also showed an increase in growth rate in the absence of the streptomycin (p<0.05). The latter included nine out of the ten populations of the *ubiH* mutant, and three populations of the *selB* mutant. These results demonstrate that for both mutants, substantial improvement in fitness (2-20% for the *selB* mutant and 28-41% for the *ubiH* mutant), including compensation for the initial fitness cost, occurred during growth at sub-MIC streptomycin levels after only 250 generations of evolution (Fig. 4A and 4B).

**Fig. 4.**
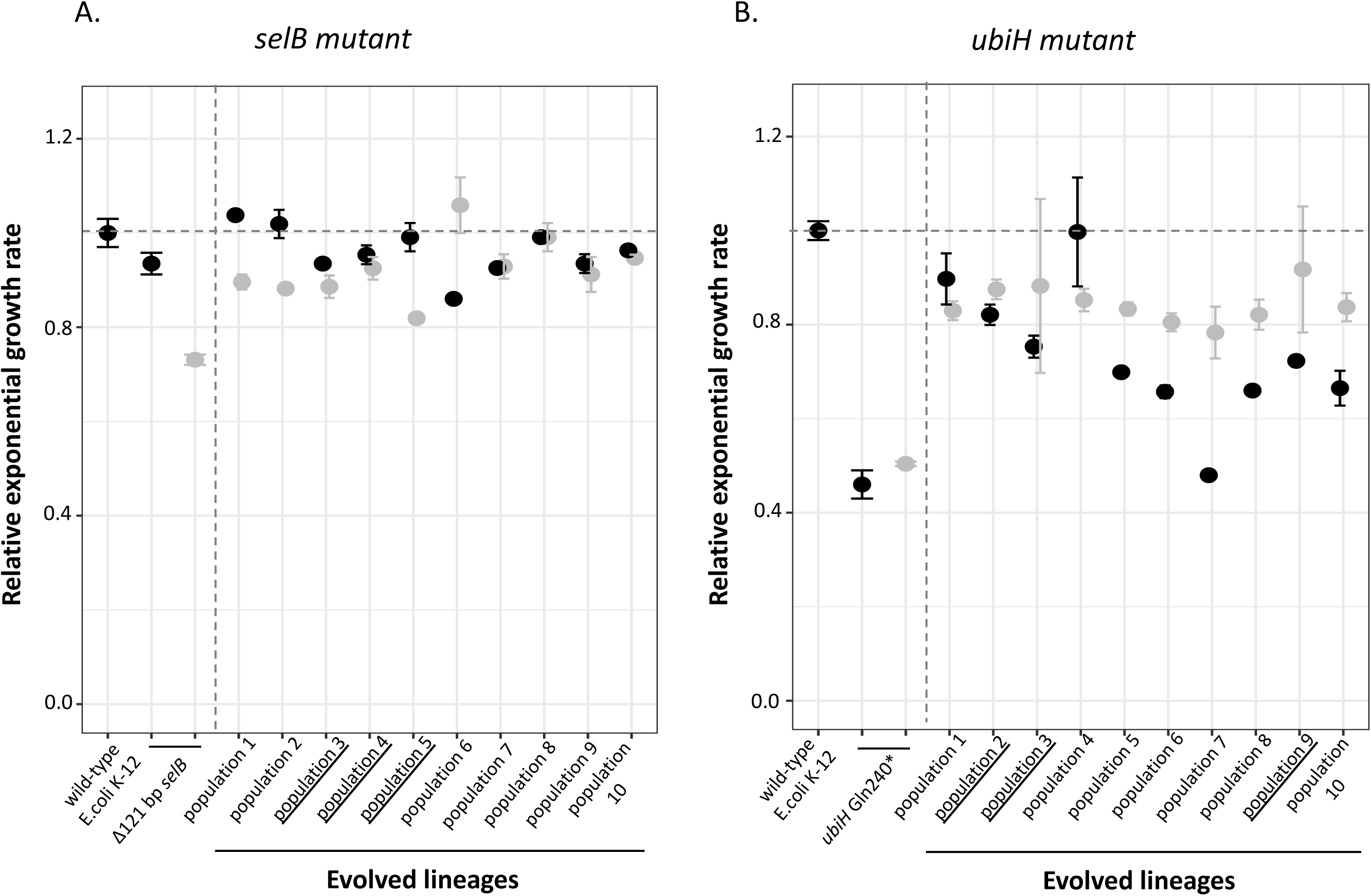
Relative exponential growth rates of populations of streptomycin resistant *E. coli selB* and *ubiH* mutants evolved at sub-MIC of streptomycin. Relative growth rates of the ancestral strains and evolved populations of the streptomycin resistant *selB* (A) and *ubiH* (B) mutants selected at streptomycin concentration of 0.8X MIC_wt_liquid_ after 250 generations are shown in the presence (grey) and absence of the antibiotic (black). Growth rates are shown relative to the growth rate of the susceptible MG1655 strain measured in the absence of the antibiotic. Three biological replicates were used in each case. Error bars show standard deviations. Individual clones were isolated from underlined populations for further characterization.

Next, clones were isolated from three populations for each mutant (underlined in Fig. 4; population 3,4 and 5 for the *selB* mutant and population 2,3 and 9 for the *ubiH* mutant) at the end point of the experiment and growth rates were measured both in the presence and absence of the antibiotic (Table 2). As compared to the wild-type ancestral strain, the *selB* mutant had a fitness reduction of 7%, which further increased to 27% in the presence of streptomycin (at 8 mg/L i.e. 0.8X MIC_wt_liquid_). Two out of the three evolved clones (from population 3 and 5) of this resistant mutant showed an increase in relative exponential growth rate by 10% at streptomycin concentrations of 8 mg/L (0.8X MIC_wt_liquid_), whereas in the absence of streptomycin they showed a smaller ∼ 3% reduction in relative growth (Fig. 5A). The third evolved clone (from population 4) showed a complete compensation of growth rate to that of the wild-type strain, both in the presence and absence of the antibiotic (Fig. 5A). The resistant mutant with the *ubiH* mutation had an initial fitness cost of 50% both with and without the antibiotic (Fig. 5B) whereas the evolved clones showed complete compensation of the growth defect as compared to the wild-type strain in the absence of the antibiotic, and a 30% increase in fitness in the presence of the antibiotic (Fig. 5B).

**Fig. 5.**
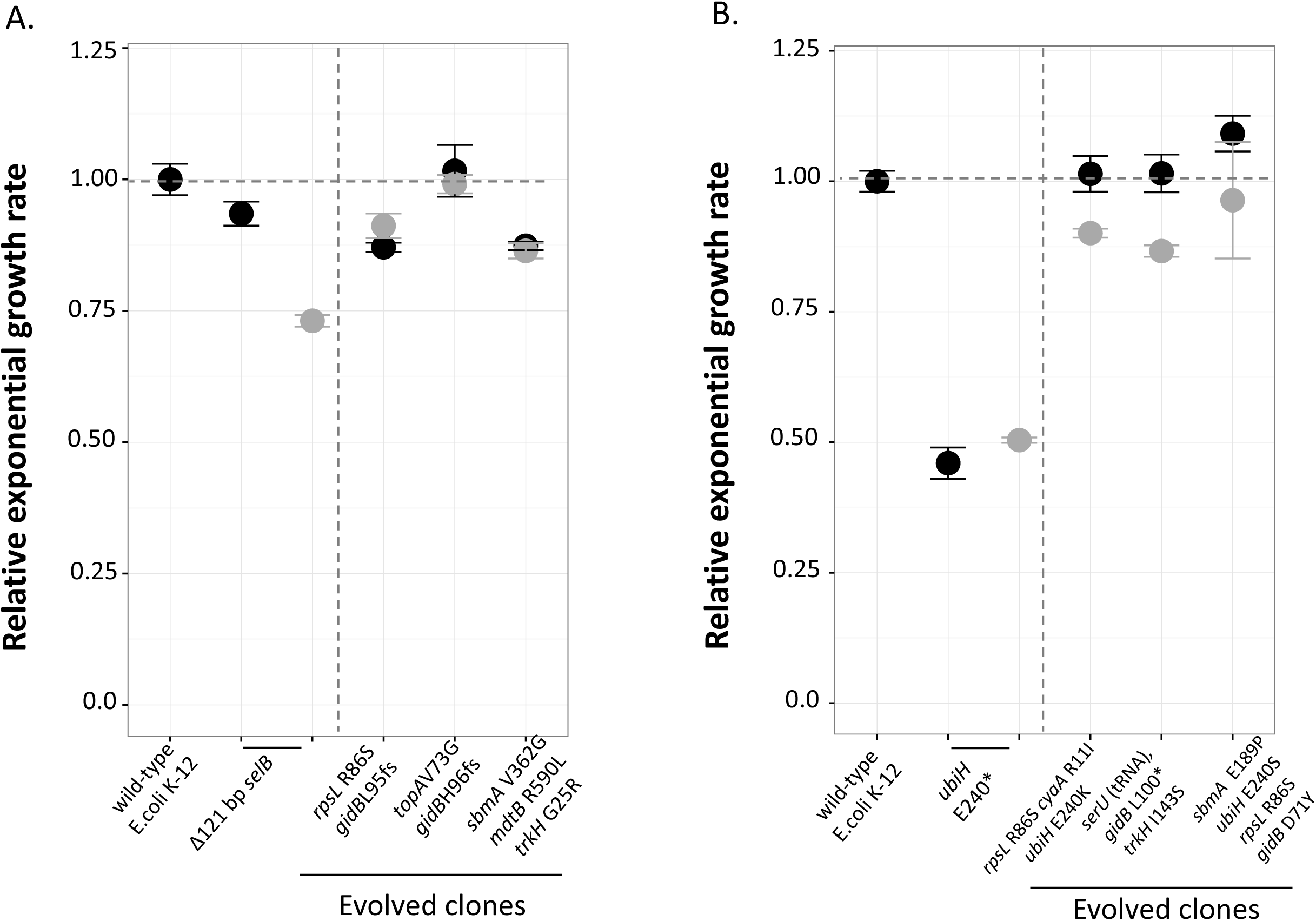
Relative exponential growth rates of evolved clones of streptomycin resistant *E. coli selB* and *ubiH* mutants evolved at sub-MIC levels of streptomycin. Relative growth rates of the ancestral strain and evolved clones of streptomycin resistant *selB* (A) and *ubiH* (B) mutants selected at a streptomycin concentration of 8 mg/L (0.8X MIC_wt_liquid_) after 250 generations, measured in absence of antibiotic (black) and in the presence (grey) of streptomycin concentrations of 0.8X MIC_wt_liquid_. Growth rates are shown relative to growth rate of susceptible MG1655 strain measured in the absence of the antibiotic. Three biological replicates were used in each case. Error bars show standard deviations.

**Table 2:**
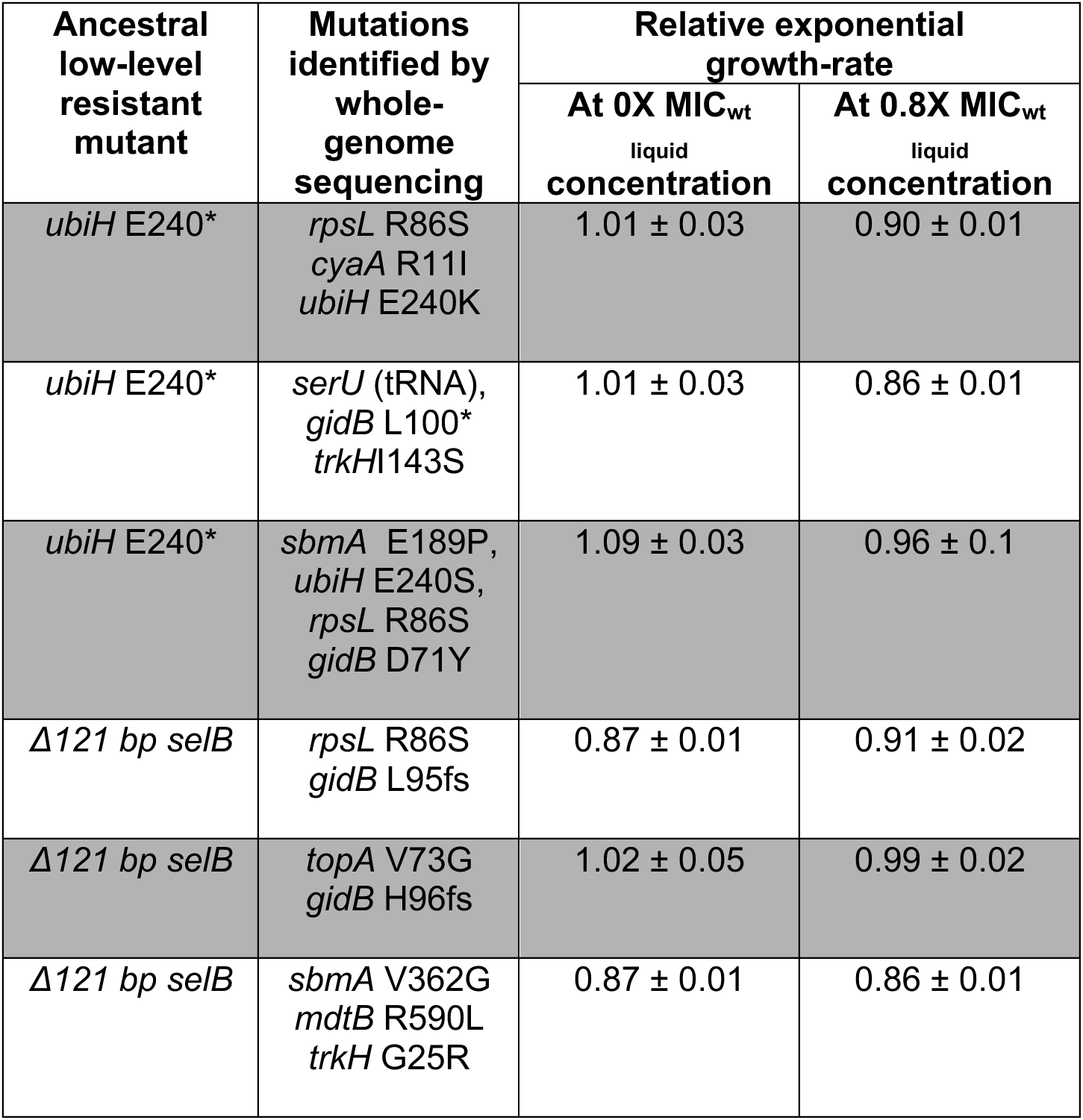
Clones of low-level resistant *E. coli* mutants *ubiH* E240* (premature stop codon at position 240) and Δ121 bp *selB* isolated after 250 generations of evolution at streptomycin concentrations of 0.8X MIC_wt**_**liquid_. Growth rates are shown relative to the growth rate of susceptible MG1655 strain measured in absence of antibiotic. All amino acids are denoted as single-letter codes A * denotes a stop-codon, and a fs denotes a frame-shift mutation.

| Ancestral low-level resistant mutant | Mutations identified by whole-genome sequencing | Relative exponential growth-rate |  |
| --- | --- | --- | --- |
|  |  | At 0X MIC<sub>wt</sub><br>liquid concentration | At 0.8X MIC<sub>wt</sub><br>liquid concentration |
| *ubiH* E240* | *rpsL* R86S<br>*cyaA* R11I<br>*ubiH* E240K | 1.01 ± 0.03 | 0.90 ± 0.01 |
| *ubiH* E240* | *serU* (tRNA),<br>*gidB* L100*<br>*trkH* I143S | 1.01 ± 0.03 | 0.86 ± 0.01 |
| *ubiH* E240* | *sbmA* E189P,<br>*ubiH* E240S,<br>*rpsL* R86S<br>*gidB* D71Y | 1.09 ± 0.03 | 0.96 ± 0.1 |
| $\Delta 121$ bp *selB* | *rpsL* R86S<br>*gidB* L95fs | 0.87 ± 0.01 | 0.91 ± 0.02 |
| $\Delta 121$ bp *selB* | *topA* V73G<br>*gidB* H96fs | 1.02 ± 0.05 | 0.99 ± 0.02 |
| $\Delta 121$ bp *selB* | *sbmA* V362G<br>*mdtB* R590L<br>*trkH* G25R | 0.87 ± 0.01 | 0.86 ± 0.01 |

### Growth-rate measurements and whole-genome sequence analysis of evolved clones of *selB* and *ubiH* mutants selected at sub-MIC of streptomycin

To examine the characteristics of the adaptive response at sub-MIC of streptomycin, growth-rate measurements (at different streptomycin concentrations) and whole-genome sequence analysis was performed for individual clones isolated from three evolved populations each of *selB* and the *ubiH* mutant (underlined in Fig.4). For the *selB* mutant, mutations that are known to confer high-level streptomycin resistance were observed in two of the three clones (Fig. 5A, Table 2). Thus, the first clone had mutations in genes *rpsL* and *gidB* (Table 2), and grew at the highest concentration of streptomycin tested 100 mg/L (10X MIC_wt_liquid_, Fig. 6A) without any growth reduction at this concentration as compared to growth without antibiotic (Fig. 5A). The second clone had mutations in genes *gidB* and *topA,* and also demonstrated an increased MIC with no appreciable growth defect at the streptomycin concentrations where growth was observed i.e. at 0.8X and 3.2X MIC_wt_liquid_, however this clone did not grow at higher streptomycin concentrations of 100 mg/L i.e 10X MIC_wt_liquid_ (Fig.6A). The third clone had mutations in genes *sbmA, mdtB* and *trkH.* This clone also demonstrated an increased MIC, but the growth rate decreased with an increase in streptomycin concentrations (Fig. 6A). Out of this set of genes, *rpsL* (which encodes small subunit ribosomal protein S12) and *gidB* (which encodes a methyltransferase that is responsible for methylation of the 16S rRNA component of the ribosome) are target mutations, while *topA* (DNA topoisomerase 1), *sbmA* (peptide antibiotic transporter) and *trkH* (low-affinity potassium transport system) have been implicated in aminoglycoside resistance by influencing drug uptake (Lavina et al. 1986; Mogre et al. 2014; Wistrand-Yuen et al. 2018).

**Fig. 6.**
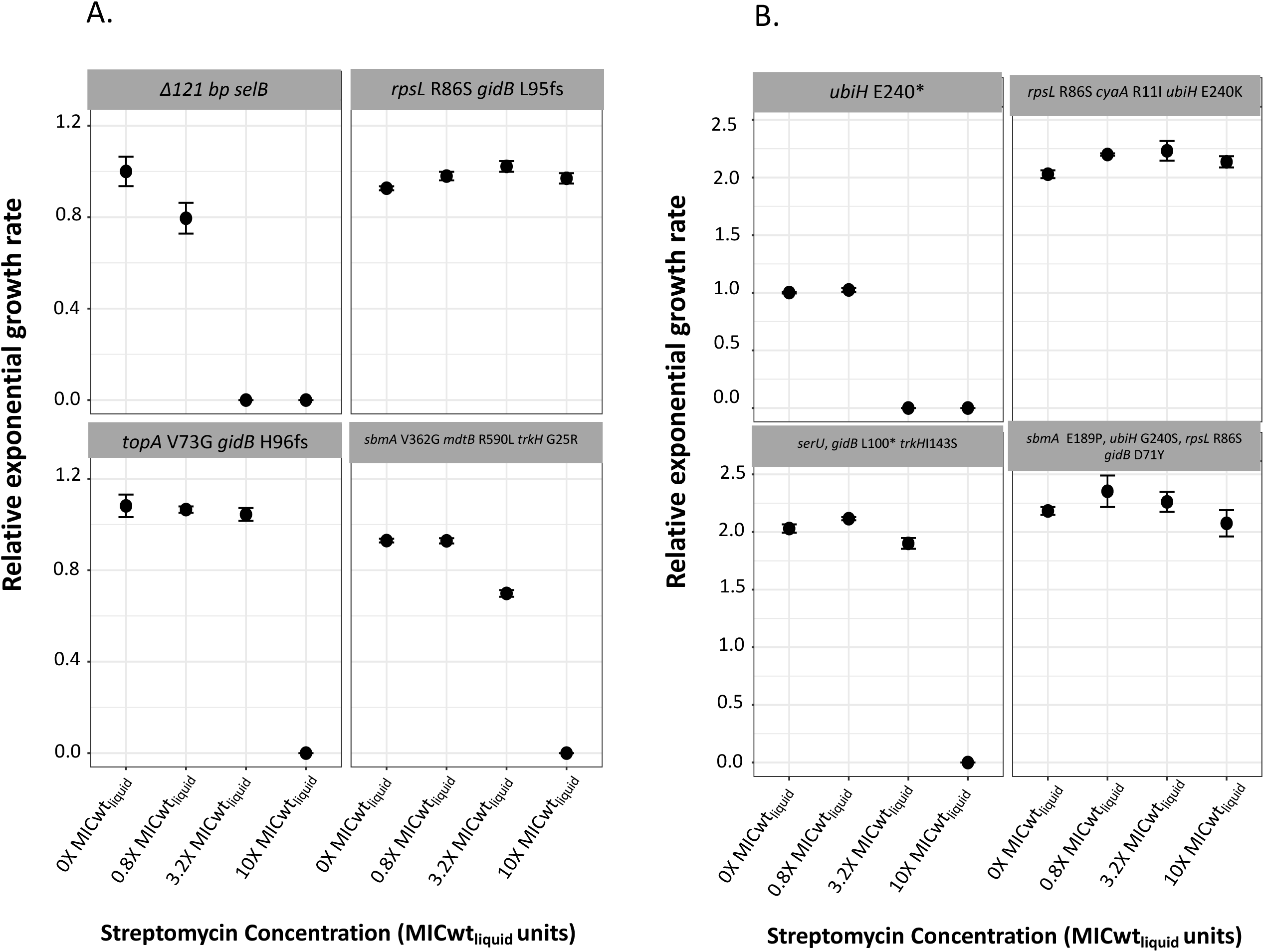
Growth-rates of evolved clones of streptomycin resistant *E. coli selB* and *ubiH* mutants at different streptomycin concentrations. Relative growth rates of evolved clones of streptomycin resistant *selB* (A) and *ubiH* (B) mutants selected at a streptomycin concentration of 0.8X MIC_wt_liquid_ were measured at different concentrations of streptomycin. Growth rates are shown relative to growth rate of mutants *ubiH* and *selB* measured in the absence of the antibiotic. Three biological replicates were used in each case. Error bars show standard deviations.

Evolved clones of the *ubiH* mutant also had mutations in target genes (*rpsL* and *gidB*) and in genes that have been implicated in streptomycin resistance i.e. *sbmA* (Mogre et al. 2014) and *cyaA* (adenylate cyclase, Table 2). Additionally, in each of these three evolved clones, mutations were observed that removed the fitness cost by changing the premature stop codon of the original mutation (at position 240 in the *ubiH* gene) to a missense mutation (Table 2, Fig 5B). Thus, in two cases the mutation changed a stop codon to a sense codon (stop240K and stop240S) whereas in the third case a suppressor mutation in the *serU* tRNA was observed (position 35 G to T, changing the sequence that codes for the anticodon from CGA to CTA), allowing suppression of the premature stop by insertion of the amino acid serine. Two of the evolved clones with *rpsL* mutations grew over the complete range of streptomycin concentrations tested, while the third mutant that had mutations in genes *gidB*, *serU* and *trkH* also had an increased MIC; additionally we did not observe any appreciable growth defect with increasing streptomycin concentrations for any of these mutants (Fig. 6B).

### Enrichment of higher-level resistant mutants in evolved populations of both the *selB* and *ubiH* mutants

We further characterized the adaptation of each population to the sub-MIC selective regime by measuring the frequency of high-level resistant mutants at the end-point of the evolution experiment. Populations were screened for resistant mutants at streptomycin concentrations of 16X MIC_wt_solid_ (64 mg/L) and 25X MIC_wt_solid_ (100 mg/L). For the *selB* mutant, three out of the ten evolved populations (populations 3,6 and 7) showed fixation of high-level resistant mutants growing at 25X MIC_wt_solid_, while in a fourth case (population 10), high-level resistant mutants at 25X MIC_wt_solid_ were observed at a frequency of ∼10^-5^ (Fig. 7A). Three other populations exhibited low frequencies (∼10^-6^) of mutants that could grow at 16X MIC_wt_solid_ (Fig. 7A, populations 4,5 and 8). For the remaining three populations (population 1,2 and 9), high-level resistant mutants were not observed.

**Fig. 7.**
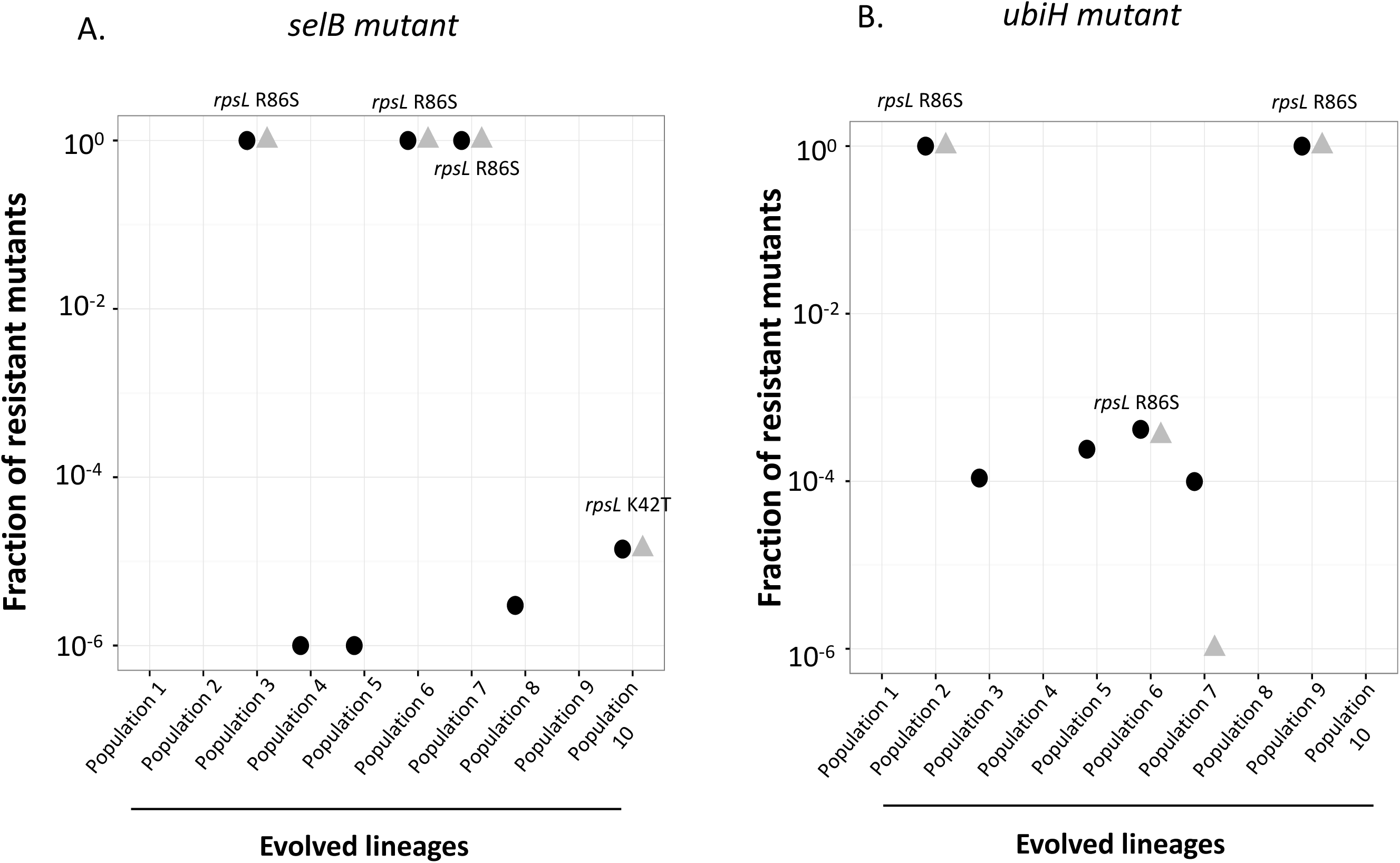
Enrichment of high-level resistant *E. coli* mutants in populations evolved at sub-MIC of streptomycin. Fraction of the total population comprised of high-level resistant mutants was measured in evolved populations of streptomycin resistant *selB* (A) and *ubiH* (B) mutants after 250 generations of growth at streptomycin concentration of 0.8X MIC_wt_liquid_. High-level mutants were selected at concentrations of 16X MIC_wt_solid_ (indicated as black circles) and 25X MIC_wt_solid_ (indicated as grey triangles). Y-axis is plotted on a log-scale.

Similar trends were observed for the evolved populations of the *ubiH* mutant (Fig. 7B). For two populations (population 2 and 9), high-level resistant mutants growing at 25X MIC_wt_solid_ had reached fixation, while in two other populations (population 6 and 7) these had reached a frequency of 10^-6^ to 10^-4^. In three populations (population 3,5 and 7), high-level resistant mutants growing at 16X MIC_wt_solid_ were observed at frequencies varying between 10^-6^ to 10^-4^; this included population 7 that also showed enrichment for high-level resistant mutant growing at 25X MIC_wt_solid_. The remaining four populations did not show any enrichment of high-level resistance mutants (population 1,4,8 and 10).

### Prevalence of *rpsL* R86S mutation in high-level resistance mutants enriched in the evolving populations

One unexpected observation in our experiments was the high prevalence of the *rpsL* R86S mutation in the high-level resistant mutants selected in evolving populations of the *selB* and *ubiH* mutants. Three of the six mutants that were whole-genome sequenced contained this mutation. To further investigate the prevalence of this *rpsL* variant, we isolated high-level resistant mutants from the twenty populations (ten populations for each low-level mutant isolated on 25X MIC_wt_solid_), and sequenced the *rpsL* gene. Out of eight different populations (populations 3,6,7 and 10 for the *selB* mutant and populations 2,6,7 and 9 for the *ubiH* mutant), six contained the *rpsL* R86S mutation, one had the *rpsL* K42T mutation, and one had no mutation in the *rpsL* gene (Fig. 7). To determine the reason for the prevalence of this variant, we constructed different known *rpsL* variants and compared their respective pharmacodynamic curves (Fig.8). Although the *rpsL* R86S variant is cost-free in the absence of the antibiotic, it either had similar or lower growth rates in comparison to other *rpsL* variants at the streptomycin levels at which the sub-MIC evolution was performed. Interestingly the *rpsL* R86S variant also had the lowest MIC (3.2X MIC_wt_liquid_) compared to the other *rpsL* variants. Thus, the data on the growth rate and the resistance levels of the *rpsL* R86S mutant itself cannot explain the prevalence of this variant in our experiments, suggesting potential epistatic interactions between the low-level resistant mutations (*ubiH* and *selB*) and the *rpsL* R86S mutation. We tried to address this hypothesis by constructing strains carrying different combinations of *rpsL* alleles and the *ubiH* or *selB* mutations but rapid accumulation of off-target mutations during strain construction precluded this possibility.

**Fig. 8.**
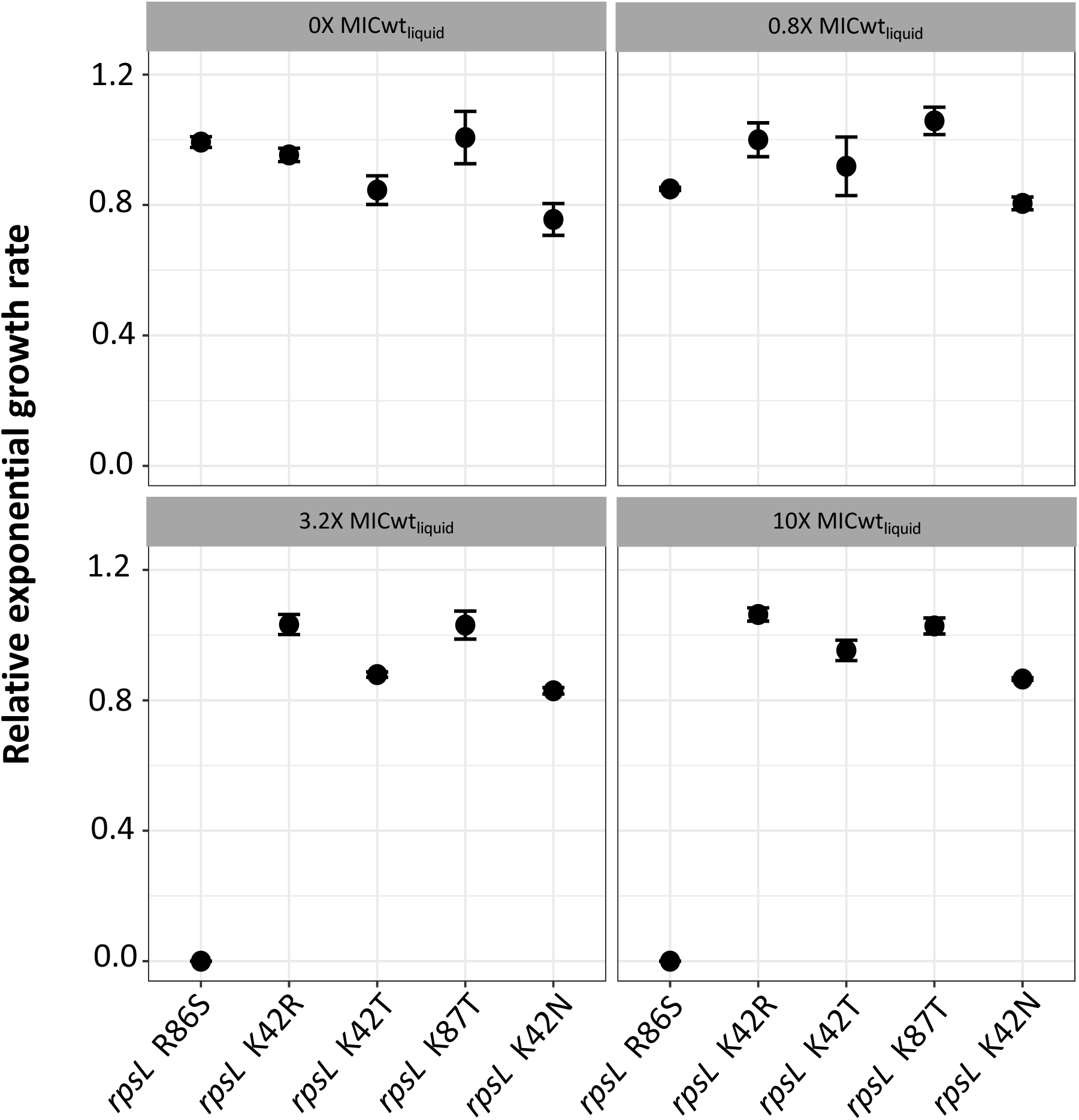
Growth-rates of streptomycin resistant *E. coli rpsL* mutants at different streptomycin concentrations. Relative growth rates of the different *rpsL* mutants were measured at different concentrations of streptomycin. Growth rates are shown relative to growth rate of a susceptible MG1655 strain measured in the absence of the antibiotic. Three biological replicates were used in each case. Error bars show standard deviations.

### Maintenance of the slow-growing *ubiH* mutant in susceptible population at sub-MIC of streptomycin

The maintenance of resistant mutants in a susceptible microbial population at sub-MIC of an antibiotic is expected to depend on the cost of the resistance mutation versus the selective effect of the antibiotic (Gullberg et al. 2011; Gullberg et al. 2014). To investigate this idea for the *ubiH* mutant, competition experiments were performed starting with equal proportions of the susceptible wild type *E. coli* strain and the slow growing *ubiH* mutant in the presence (streptomycin concentration 0.25X MIC_wt_liquid_) and absence of streptomycin (Fig. 9A). Both strains were genetically tagged with genes encoding fluorescence proteins allowing us to determine the change of frequency of each strain over the course of the experiment. As a control, the susceptible strain was also marked with the different fluorescent markers and competed in the presence and absence of the antibiotic for the same period of time to determine the cost of the fluorescent proteins (Fig. 9B). These experiments showed that the *ubiH* mutant is outcompeted by the susceptible strain in the absence of the antibiotic in less than 50 generations, but maintained in the population in the presence of sub-MIC of streptomycin during the experiment. Although in most populations the *ubiH* mutant was reduced in frequency as compared to susceptible ancestral *E. coli* strain, the rate at which this occurred was much slower than in the absence of the antibiotic (Fig. 9A). In at least two out of the eight populations the reduction in mutant frequency was much slower as compared to the remaining populations, indicating the occurrence of compensatory mutations (Fig. 9A). Thus, these results demonstrate that costly resistant mutants can be maintained in the population with susceptible strains for longer periods of time which then could allow them to acquire compensatory and/or resistance mutations.

**Fig. 9.**
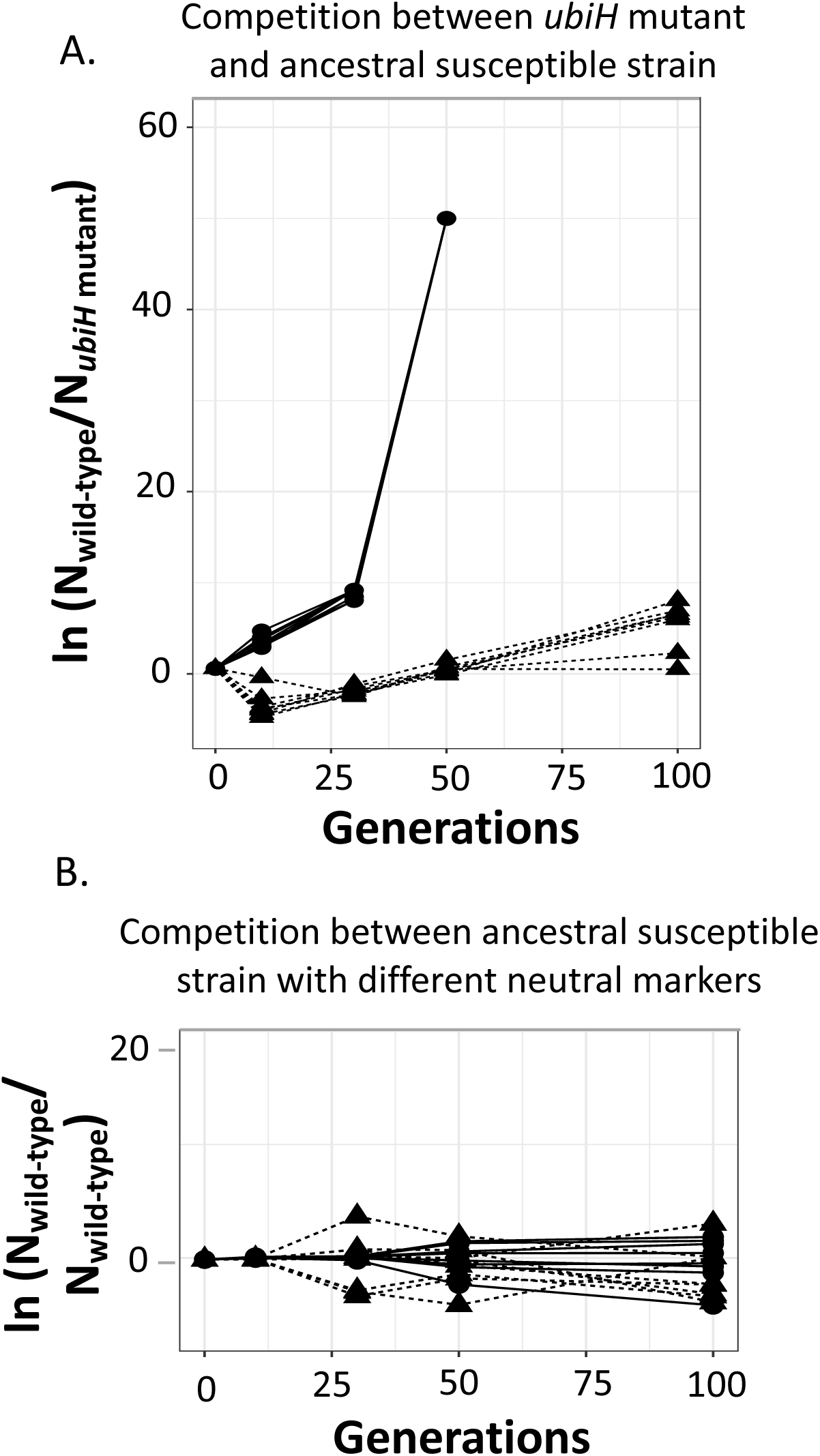
Competition experiments between a slow-growing *ubiH* mutant and the susceptible ancestral *E. coli* strain at sub-MIC of streptomycin. Competition experiments were performed between the *ubiH* mutant and the susceptible ancestral strain (A), and between the susceptible ancestral strain containing neutral fluorescent markers (B) in the absence (solid lines with circles) and presence (dashed lines with triangles) of streptomycin (0.25X MIC_wt_liquid_). In each case, the strains were fluorescently marked and the frequency was measured at five different time points (0, 13, 30, 50 and 100 generations) for eight biological replicates.

## DISCUSSION

The main objective of this study was to determine how antibiotic resistance and bacterial fitness evolve during growth in presence of sub-MIC antibiotic concentrations. To this end, two low-level streptomycin resistant mutants were chosen (MIC of streptomycin for each mutant was 3.2X MIC_wt_liquid_), for evolution experiments carried out at sub-MIC of streptomycin. The choice of resistant mutants was based on the observation that they had identical MICs but differences in fitness costs, and in how growth of these mutants varied at different concentrations of streptomycin (Fig. 3). Thus, in the absence of the drug the *selB* mutant had a small reduction in growth-rate while the *ubiH* mutant had a large reduction in growth-rate. Despite these differences, we observed that compensatory evolution and evolution for increased antibiotic resistance was common for both mutants under sub-MIC selection regimes, though compensation was more prevalent for the *ubiH* mutant.

Whole-genome sequencing of the evolved clones showed that compensation of the physiological cost of the *ubiH* E240* mutation was due to selection of specific mutations that allowed the expression of the UbiH protein, i.e. by restoration of the original function. However, in each of these clones we also observed mutations that are known to increase resistance to streptomycin. On the other hand, all the mutations observed in the evolved clone of the *selB* mutant conferred increased streptomycin resistance. Thus, in this case the increase in fitness was predominantly due to the clones becoming more resistant, and not due to the compensation of the physiological cost conferred by the starting resistance mutation. Results similar to this latter scenario was also observed in another study where selection for increased fitness in fluoroquinolone resistant mutants in the absence of the antibiotic also resulted in an increase in fluoroquinolone resistance (Marcusson et al. 2009).

Overall, our study demonstrates that depending on the cost of the resistance mutations, compensatory evolution and resistance evolution could interact to determine the evolutionary response in populations evolving at sub-MIC antibiotic levels. Compensatory mutations allow maintenance of costly resistant mutants in the population, as has been demonstrated when no antibiotic is present (Andersson and Hughes 2010; Andersson and Hughes 2011; Moura de Sousa et al. 2015), as well as under weak selection for antibiotic resistance (sub-MIC) (Westhoff et al. 2017). Our results add to these earlier studies and show that high-cost, low-level resistant mutants can be maintained in the population with the susceptible wild-type strain at sub-MIC antibiotic levels for a longer time, allowing them to accumulate compensatory mutations.

Another major finding of our study is that the “mutational space” for streptomycin resistance (i.e. all mutations that confer a significant increase in MIC of streptomycin) is much larger than previously thought. This is important in terms of evolution at sub-MIC of antibiotics, since given the high mutation rate to low-level resistance (>10^-7^/cell/generation) these high frequency, low-level resistant mutants could serve as stepping-stones for evolution of higher-level resistance via rarer mutations. This could occur by allowing bacterial survival and slow growth in the presence of the antibiotic, increasing the probability for higher-level resistance mutations to emerge. In our study, apart from the previously described aminoglycoside-modifying enzymes and target modification mutations in the *rrn*, *rpsL* and *gidB* genes, mutations in several other genes were shown to confer low- or high-level resistance, either singly or in combination ((Wistrand-Yuen, et al. 2018) and results presented here). For example, mutations in *cydA/mngB, cydB, cyoB, dxs, hemC, ispH, ndh, nuoG, selB, sst, trkH, ubiA, ubiH, ubiI, ygiC, yfjG* and *znuA* genes have been implicated in streptomycin resistance. Interestingly, in several of the *ubiH* mutants where the stop codon was reverted to a sense codon several new mutations conferring resistance replaced the original *ubiH* mutation (Table 2). Finally, it is notable that a substantial number of the target genes encode proteins that are associated with the membrane and the electron transport chain and a hypothesis for their action, which has some experimental support, is that the mutations reduce the proton motive force that is required for streptomycin uptake (Lázár et al. 2013; Wistrand-Yuen et al. 2018).

---

**Supplementary figure.1.**
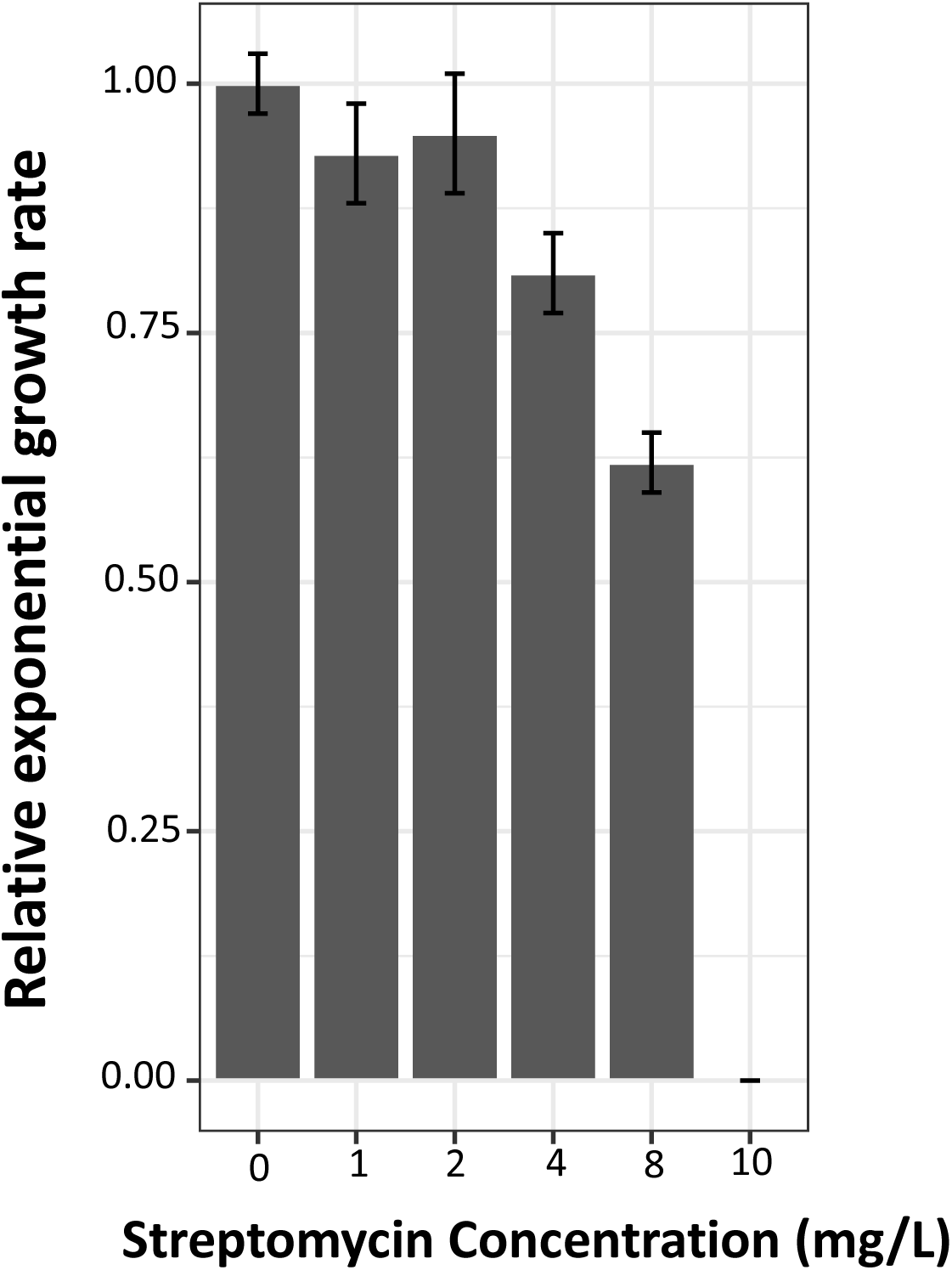
Dosage response curve for the susceptible *E.coli* MG1655 strain. All values are plotted relative to the growth-rate of the susceptible wild-type strain in the absence of streptomycin.

